# Informing antiviral effectiveness for influenza A and SARS-CoV-2 by quantifying within-host interaction between transmission and immunity

**DOI:** 10.1101/2025.03.11.642706

**Authors:** Deven V. Gokhale, Miria Ferreira Criado, Dawne K. Rowe, S. Mark Tompkins, Pejman Rohani

## Abstract

Antiviral therapies are among the most effective pharmaceutical interventions in treatment of a variety of viral pathogens. To optimize the antiviral effectiveness it is crucial to characterize the relationship between multiple cellular modes of antiviral action and the complex response of the host’s innate immune system relative to the within host dynamics of a proliferating virus. Since their introduction in 1968 and 2019, Influenza A virus (IAV) H3N2 and Severe Acute Respiratory Syndrome Coronavirus-2 (SARS-CoV-2), respectively, have caused unprecedented damage on the public health infrastructure globally. In addition to the substantial burden of morbidity and mortality around the world, both viruses have the potential of undergoing evolution leading to antigenic escape from the prevailing interventions. These biological characteristics advocate for urgent development of effective antivirals for the treatment of IAV and SARS-CoV-2.

In this multi-stage study, we develop a suite of within-host models encompassing a number of hypotheses regarding virus-specific innate host functional responses and their impacts on the proliferation of IAV H3N2 and SARS-CoV-2 viruses. We use likelihood-based statistical inference to confront these hypotheses with infection data on IAV H3N2 and SARS-CoV-2 from infection experiments in ferrets. Upon identifying the best-fitting model of within-host dynamics, we can quantify the potential impact of antiviral drug therapy as a function of effectiveness and timing of initiation.

We find significant mechanistic differences between the infection dynamics of H3N2 IAV and SARS-CoV-2 and associated model parameters. The treatment consequences of these differences are that SARS-CoV-2 is harder to control with antivirals, requiring earlier initiation and a more effective drug.

**Author summary:** Antiviral drugs are prophylactic chemical agents that are used to contain several viral infections. Influenza A H3N2 virus (H3N2) and Severe Acute Respiratory Syndrome Corona Virus-2 (SARS-CoV-2) are very important viral pathogens that have caused unprecedented, global public health damage in the recent times. This makes development of antiviral drugs crucial along with other pharmaceutical prophylactics like vaccination. To optimize the pathogen specific effectiveness, however, it is necessary to simultaneously explore the relationship among the intra-host viral kinetics, immune dynamics and modes of antiviral action. In this article we theoretically analyze antiviral action of a drug in union with effect of the host’s innate immune system in containing infections of SARS-CoV-2 and H3N2 in infection experiments with ferrets. We find fundamental differences in the requisite antiviral effectiveness which, we posit, is due to substantially different inter-cellular proliferation potential between the two viruses.

## Introduction

Antiviral therapies are among the most effective pharmaceutical interventions in the control and treatment of a variety of viruses [1]. These drugs function by interrupting the viral life cycle within host cells, with the ultimate aim of preventing infection, shortening the infectious period and curtailing the chain of transmission [2]. In addition to these direct effects, antivirals have been proposed as a candidate public health response both to contain outbreaks [3, 4], and as a strategy for prophylactic prevention of zoonotic spillover [5]. The efficacy and prescribed usage of any particular host-targeting antiviral depends on the cellular pathway that it manipulates in addition to the target viral kinetics [6, 7].

The infection process for respiratory viral infections begins with inhalation of aerosolized droplets emanating from an infectious host [8]. Disseminated viral particles then enter the upper respiratory tract and following a rapid replication phase in the mucosal epithelial cells, proliferate throughout the respiratory system. Viral production triggers a multifaceted cascade of rapid innate immune responses aiming for efficient control of the infection [9]. In devising antivirals, therefore, it is crucial to characterize the timing, mechanism and magnitude of the host’s functional immune response relative to the virus replication cycle and proliferation [2, 10].

Animal infection experiments can be very useful in identifying the operative pathways and titrating the potential effectiveness of candidate drugs [11–14]. There are several animal models available to assess infection, disease, transmission, and/or efficacy of interventions against influenza A virus and SARS-CoV-2, with each infection model having specific strengths and weaknesses. For influenza, ferrets and mice have been used since the first isolation of human influenza in the 1930s and dominate the experimental infection field (recently reviewed in [12]). Ferrets are naturally susceptible to H3N2 infection and efficient transmitters to naïve ferrets [12, 15]. The ferret model is hampered by a lack of reagents, however this drawback is slowly changing [12, 13]. Animal models for SARS-CoV-2 infection were rapidly explored and developed over the past 18 months. Like influenza, ferrets were quickly assessed as models of infection, disease and transmission (recently reviewed in [14]). As in the case for H3N2 infection, the ferret is naturally susceptible to SARS-CoV-2 infection and can transmit to naive animals via contact or aerosol [16]. The various animal infection model allied with mechanistic computational approaches can permit the testing of multiple hypotheses regarding viral proliferation and innate immunity. The predictions of such models can then be validated by subsequent concise *in vivo* infection studies.

Directly transmissible respiratory viruses like the influenza A H3N2 virus (H3N2), historically, and Severe Acute Respiratory Syndrome Coronavirus-2 (SARS-CoV-2), more recently, have exacted a heavy toll on public health globally. Since its introduction into the human population following the 1968 influenza pandemic, H3N2 has accounted for substantial annual deaths globally [8]. The estimated global disease burden for severe seasonal influenza cases is around 3-5 million cases annually with a median of 400,0000 recorded deaths [17, 18], with many of these attributed to H3N2. At the time of writing, the SARS-CoV-2 pandemic has already claimed over 5 million lives worldwide [19], inflicting an unprecedented strain on global systems, and especially healthcare [20, 21]. In addition to their substantial burden on morbidity and mortality, RNA viruses such as SARS-CoV-2 and H3N2 pose additional threats because of their potential for rapid divergent evolution [22]. Examples include the recent emergence of the highly transmissible B.1.617.2 (delta) and B.1.1.529 (omicorn) variants of SARS-CoV-2 [23, 24] and the repeated punctuated emergence of novel antigenic clusters of influenza viruses [25–27]. The combination of efficient transmission of these viruses, their potential for antigenic escape and the toll they exact on our population makes the potential use of antiviral therapies for their rapid and efficient control especially timely, along with other critical pharmaceutical interventions such as vaccination.

In this article, we examine the consequences for antiviral treatment on within-host dynamics of SARS-CoV-2 and H3N2 using a combination of data from animal infection experiments, mathematical models, and statistical inference. In the first phase, we present a suite of mechanistic models of varying degrees of complexity that aim to elucidate the potential role of different components of the host innate immune system during an infection. By comparing our best-fitting models, we note major differences in the relative contribution of the host immune system in modulating the infections of H3N2 and SARS-CoV-2, with stark differences in the within-host proliferation of the two viruses. Next, to understand the impact of antiviral timing, dose and mode of action, we incorporate antiviral dynamics into our fitted within-host model. Our numerical and analytical findings indicate efficient control requires virus-specific antiviral initiation time and effectiveness.

## Materials and methods

### Infection Experiments

#### Procedure

Five to six-month-old, male and female (range 800-1200g) fitch ferrets (*Mustela putorius furo*, n=4 per group) were housed in pairs in the animal biosafety level 3 enhanced (ABSL3E) facilities at the University of Georgia. After a period of acclimation, ferrets were anesthetized with isoflurane and intranasally inoculated with H3N2, A/Singapore/INFIMH-16-0019/2016 strain (IRR FR-1590; SG/16) or SARS-CoV-2 virus (USA-WA1/2020, BEI Resources, NR-52281) diluted in 1.0 ml of Phosphate Buffered Saline (PBS) solution. Virus inoculum dose delivered to each ferret was 10^6^ PFU (plaque-forming unit)/ml of IAV H3N2 or 10^6^ FFU (focus forming units)/ml of SARS-CoV-2. Following inoculation, animals were monitored daily for clinical signs, including nasal discharge, sneezing, diarrhea, lethargy, increased respiratory rate and effort (congestion), cyanosis, neurological changes, and altered responses to external stimuli. Nasal washes, weights and temperatures were obtained every other day under light sedation. For nasal washes collection, ferrets were anesthetized with isoflurane and their nares irrigated with 3.0 ml of PBS. Infectious viruses were titrated in either Madin-Darby canine kidney (MDCK) or Vero E6 cells by PFU or FFU assays, respectively. A separate aliquot of nasal wash was processed for RNA isolation and subsequent quantification by qRT-PCR [28]. At the end of the study, infected animals were humanely euthanized.

#### Ethics Statement

All animal experiments were approved by the University of Georgia Institutional Animal Care and Use Committee (protocol number A2020 03-016). Animals were housed and maintained in accordance with the US Animal Act PL 99-158 and in accordance with guidelines stated in the “Guide for the Care and Use of Laboratory Animals”.

### Within-Host Dynamics

#### Model of Infection and Innate Immunity

To systematically explore potential interactions between viral proliferation and the host innate immune responses, we developed a family of mechanistic models incorporating a number of distinct immuno-ecological hypotheses, as illustrated in Fig. 1.

**Fig 1.**
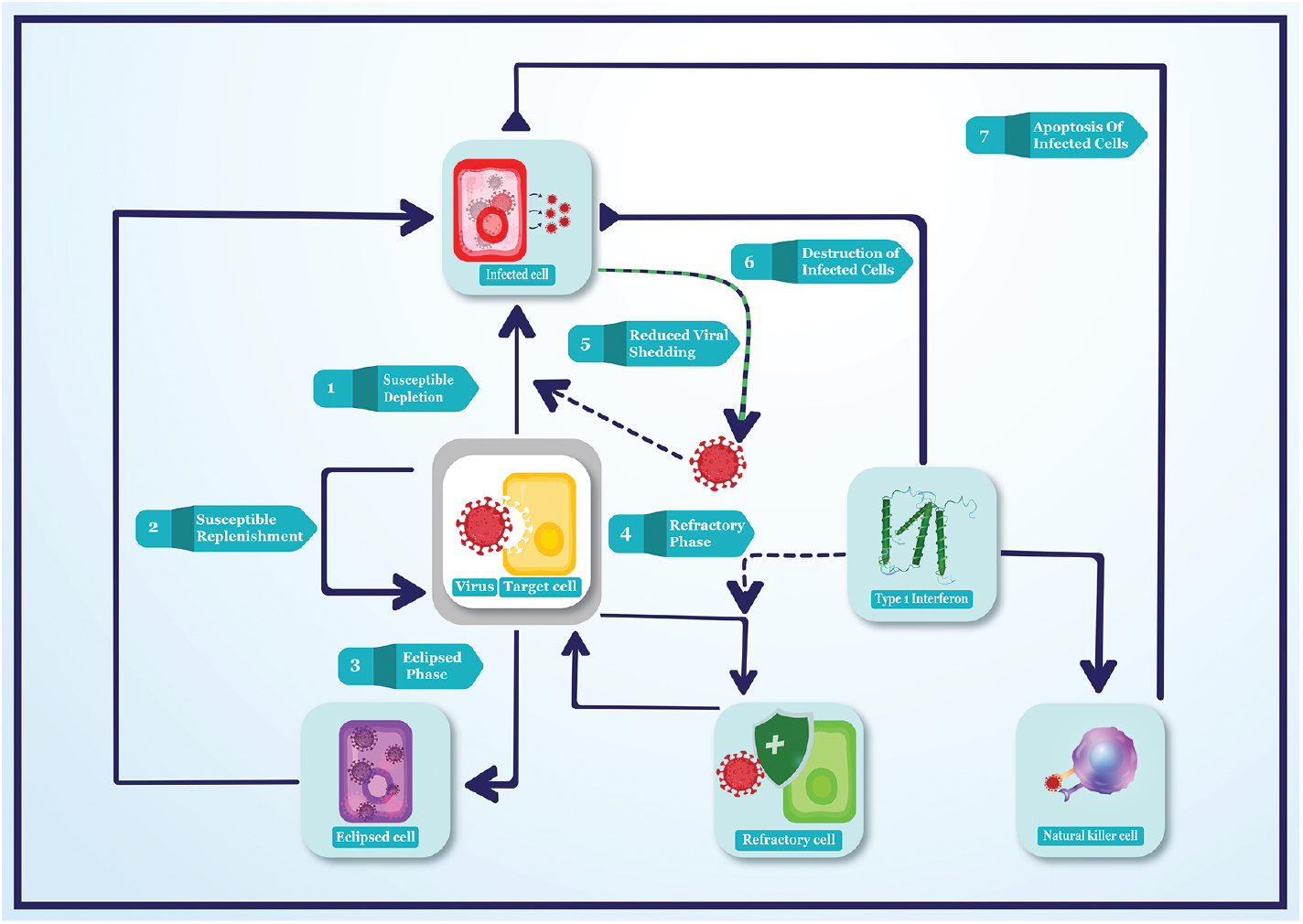
Schematic of the focal hypotheses considered in this study. All sub-populations involved in the host-virus interactions are colour coded. Arrows indicate promotion and flat-headed arrows indicate inhibition reactions describing cellular interactions.

Respiratory viruses such as H3N2 and SARS-CoV-2 commonly replicate in the epithelial cells of the upper respiratory tract [29], on target cells. In general, upon infection, cells can go through a transient eclipse phase [30] during which infectious virus is absent as virus genome undergoes intracellular replication and viral proteins are produced. Subsequently, viral particles are assembled within the infected cells, leading to the viral burst of infectious virions.

We assume target cells (denoted by *T*) are infected at the rate *β*_*T*_ and become productively infectious after a transient latency (the eclipse phase), lasting 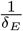 days. To be consistent with our experimental data sources, we assume nasal washes are composed of a mixture of viral RNA and viable virus. Specifically, we assume productively infectious cells (*I*) produce viral RNA (*P*) at rate *ν* and a subset (*c*) of this population is assembled into viable virus particles (*V*).

Given the short temporal duration of these experiments, typically lasting no more than 8 days, we limit our investigation around the role of the host innate immune response. Particularly, we focus on examining the action of rapidly acting, cytokine-mediated immune responses such as type I interferons *α, β* (IFN-1) and type III interferon *λ* (IFN-3) [31–33]. This family of signalling molecules generally forms the majority of the host’s initial broad sense response [9].

Three functions that are jointly associated with the release of IFN-1 and the IFN-III include

- **Increase in the prevalence of refractory epithelial cells**. Refractory cells can prevent viral entry. A high proportion of such cells can thus reduce the effective proliferation rate of the virus [34].
- **Inhibited viral shedding**. Cytokine production can promote hostile local conditions by enhancing environmental toxicity, inflammation or by recruiting resident macrophages that can attack extra-cellular viral particles. Depleting the extra-cellular viral population effectively reduces the viral production rate [35, 36].
- **Natural killer (NK) cell recruitment**. NK cells are cytotoxic lymphocytes that do not require activation to kill cells [37]. IFN-1 production can enhance the recruitment of lymphocytes such as natural killer cells, which can identify infected cells and rapidly destroy them by stimulating apoptosis thereby impeding within-host viral spread.

IFN-3, *λ* has similar functions to the IFN-1, but is generally considered to be produced earlier at mucosal surfaces and less potent than the activities of interferons *α* and *β* [38].

#### Immune-independent and immune mediated infection hypotheses

To identify the most parsimonious causal mechanism consistent with the observed viral dynamics, we formulated and contested 7 distinct hypotheses, described below. Model state variables and parameter definitions are provided in tables 1 & 2.

**Table 1.** Model compartments for the model of infection and immunity.

| Compartment | Definition |
| --- | --- |
| $T$ | Target epithelial cells, invasion sites |
| $E$ | Eclipsed epithelial cells |
| $I$ | Infectious epithelial cells, replication sites |
| $V$ | Viral titer |
| $P$ | Viral RNA |
| $R$ | Refractory epithelial cells |
| $F$ | Type I ($\alpha$ & $\beta$) type III ($\lambda$) interferon |
| $N$ | Natural killer cells |

#### Hypothesis I. Susceptible cell Depletion

The simplest model incorporates epithelial target cells (*T*) that are infected by viable virions (*V*) at *per capita* rate *β*_*T*_. Upon transmission, infected cells (*I*) produce novel viral RNA (*P*) at rate *ν* and viable viral particles (*V*) at a reduced rate (*cν*, where *c <* 1). Thus, in this model, the main driver of viral dynamics is the depletion of target cells.

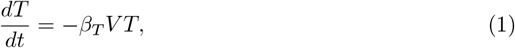

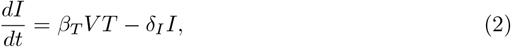

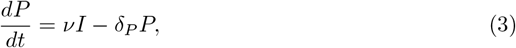

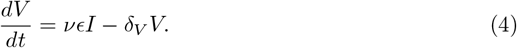

#### Hypothesis II. Susceptible Replenishment

This is an extension of the model described by equations (1)–(4) and includes the replenishment of target cells. Specifically, we assume target cell dynamics are described by logistic growth with a maximum growth rate *ν*_*T*_ and a ‘carrying capacity’ of (*κ*_*c*_).

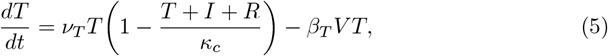

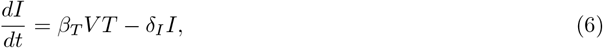

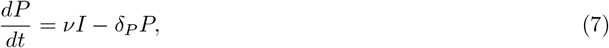

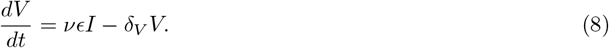

#### Hypothesis III. An eclipse phase

This model extends the system described in Hypothesis I with the inclusion of eclipsed infectious cells (*E*). The assumption here is that within infected cells, there is a period of viral replication before the cell becomes productively infectious (akin to the latent class in population models [40]). Cells in the eclipsed phase (*E*) are converted to productively infectious cells (*I*) at rate *δ*_*E*_.

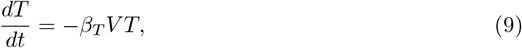

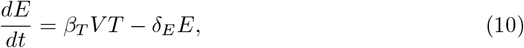

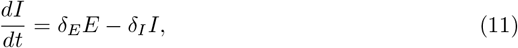

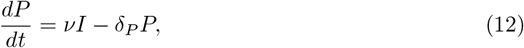

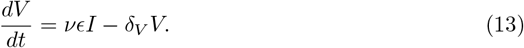

#### Hypothesis IV. IFN-induced refractory cells

Starting with the model described in equations (1)–(4), this hypothesis examines the impact of IFN-I and IFN-III (*F*) production, stimulated by the presence of infectious cells (*I*) leading to the generation of refractory target cells (*R*), which are uninfected target cells that are transiently resistant to viral attack. IFN-I and IFN-III molecules are produced at rate *ν*_*F*_ and bind to and convert target cells to refractory cells at rate *φTF*. Refractory cells are resistant to viral infection for an average of 1*/ρ* days.

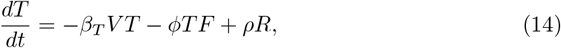

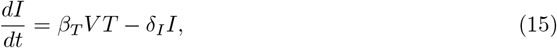

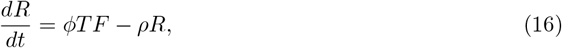

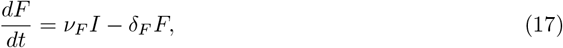

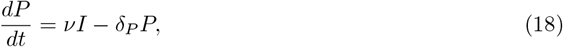

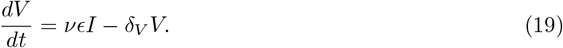

#### Hypothesis V. IFN-induced reduction in viral shedding

This model extends equations (1)–(4) by incorporating an alternative antiviral action of IFN-I and IFN-III molecules. Specifically, the release of Type I and III interferons is implicitly assumed to reduce viral shedding by an infectious cell (*I*) by a factor *e*^−*πF*^.

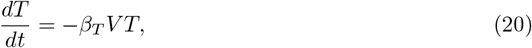

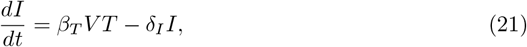

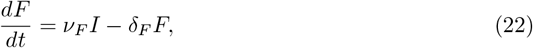

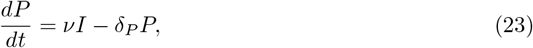

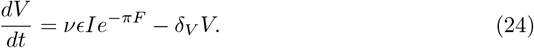

#### Hypothesis VI. Natural killer cells

This model incorporates the impact of natural killer cells (NK cells, *N*) into the default model (equations 1–4). Here, we assume IFN-I and IFN-III release stimulates the recruitment of NK cells at rate *ν*_*N*_ *F*. NK cells recruited to the site of infection kill infected cells at rate *ψNI* also leading to the clearance of the acting NK cells.

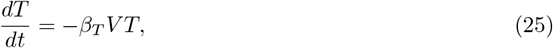

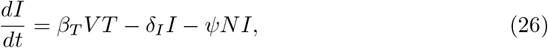

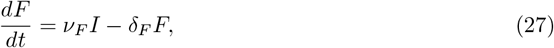

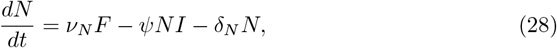

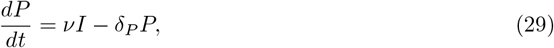

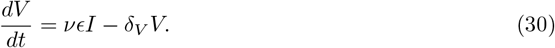

#### Hypothesis VII. Natural killer cells-2

Here, in contrast to the model presented in equations (25)–(30), we assume the release of IFN-I and IFN-III generates an inflammatory response leading to the apoptosis of infected cells (*I*) at rate *ζF*.

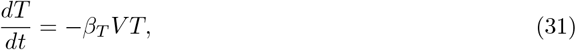

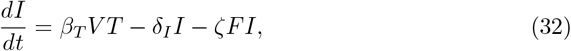

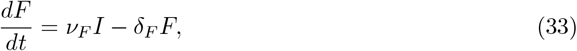

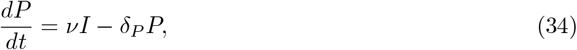

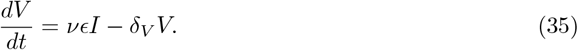

#### Derivation of *R*_0_

To summarize the within host transmissibility of a virus, we analytically derived the expression for basic reproductive number (*R*_0_) for this system. *R*_0_ for the within-host viral kinetics predicts the average number of infected cells generated by a typical infectious cell at the start of the infection [41]. We simplify the above model (Eq. 1-4) to derive the expression for (*R*_0_):

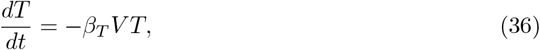

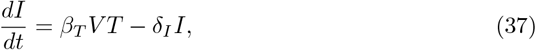

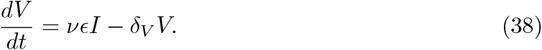

In the absence of virus, the within-host system has a stable equilibrium at (*T* ^∗^, *I*^∗^, *V* ^∗^)= (*κ*_*c*_, 0, 0). We used the next generation operator [42] to derive an expression for *R*_0_:

1. The compartments that contribute to the production of novel infections are *I* and *V*. F = [*β*_*T*_ *V T*, 0], is the vector of the rates at which new infections are generated. V = [*δ*_*I*_*I*, −*νcI* + *δ*_*V*_ *V*], is the vector of rates at which the virus flows through the system compartments.
2. For each of these vectors, F and V, we defined the Jacobian matrices, evaluated at the virus-free equilibrium. **F** is such that every element 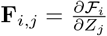, and **V** is such that every element 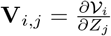, where Z = [I, V], is the vector of relevant state variables in generation of novel infected cells. This yielded 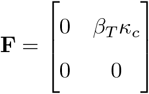 and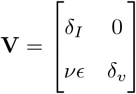.
3. The next generation operator is therefore given by 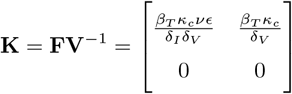. Solving the characteristic polynomial |**K** − Λ**I**| = 0 (where, **I** is the identity matrix) and identifying the dominant eigenvalue yields the expression for the basic reproductive number (*R*_0_):

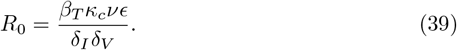

#### Sensitivity analysis

To identify model parameters critical to producing infection dynamics we conduct a sensitivity analysis on three model outcomes - (i) the peak viral load; (ii) the timing of the viral peak; and (iii) the slope of the viral decay. A sample of 500 parametric combinations is generated using a Sobol sampling scheme [43, 44] and the three model outcomes are calculated for each sampled parameter combination. The partial ranked correlation coefficient (PRCC) is calculated to quantify the association between model outcomes and individual parameter values. PRCC is an effective sampling metric to evaluate sensitivity of model outcomes on parameters [45]. We generate a PRCC bootstrapped distribution, of size 150, by sampling with replacement from the population of parameter configuration. Statistical significance of the PRCC is ascertained at 5% level of significance (LOS). Bonferroni’s correction is applied to the LOS to account for error inflation associated with multiple comparisons.

### Statistical inference and model selection

#### Observation model

For both H3N2 and SARS-CoV-2 viruses, data from ferret infection experiments is used to arbitrate among the competing models. Maximum likelihood estimation of unknown parameters is carried out by trajectory-matching in the R package *pomp* [46, 47]. *Here we assume log-normally distributed observation error. We denote our data by* 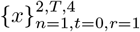, where *n* represents the data type (viable virus and RNA), *T* represents the time course of infection, and *r* represents replicate animals. The likelihood function for these data is thus given by:

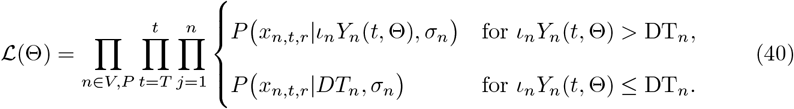

Here, *ι*_*n*_ is the data-type specific probability of under-sampling, *Y*_*n*_(*t*, Θ) is the model solution of the data-type at time *t, σ*_*n*_ is the time invariant standard deviation of the log-normal distribution whose probability density function is given as P(.|Λ_*n*_) where Λ_*n*_ is the parameter vector of this distribution (see Table 3).

**Table 2.** Model parameters with definition and units for the full model of infection and immunity. Parameters are grouped by the hypotheses and arranged in an order of increasing biological complexity.

|  | Parameter | Value/Range | Unit | Definition |
| --- | --- | --- | --- | --- |
| **Immune independent dynamics** |  |  |  |  |
| (H1)<br>Susceptible<br>depletion | $\kappa_c$ | $7 \times 10^7$ [39] | $n_T$ | Target cell population size |
| | $\nu$ | $\geq 0$ | $n_V n_T^{-1} d^{-1}$ | Viral RNA production rate |
| | $\epsilon$ | $[0,1]$ | - | Viable virus production rate |
| | $\delta_P$ | $\geq 0$ | $d^{-1}$ | Viral RNA clearance rate |
| | $\delta_V$ | $\geq 0$ | $d^{-1}$ | Viral clearance rate |
| | $\delta_I$ | $\geq 0$ | $d^{-1}$ | Infected cell clearance rate |
| | $\beta_T$ | $\geq 0$ | $n_T n_V^{-1} d^{-1}$ | Infection rate |
| (H2)<br>Susceptible<br>replenishment | $\nu_T$ | $\geq 0$ | $n_T d^{-1}$ | Target cell production rate |
| (H3)<br>Eclipsed<br>phase | $\delta_E$ | $\geq 0$ | $d^{-1}$ | Eclipse cell conversion rate |
| **Immune dynamics** |  |  |  |  |
| Type-1<br>interferon<br>dynamics | $\nu_F$ | $\geq 0$ | $d^{-1}$ | Activation rate |
| | $\delta_F$ | $\geq 0$ | $d^{-1}$ | Deactivation rate |
| (H4)<br>Refractory<br>cell<br>dynamics | $\phi$ | $\geq 0$ | $n_T n_F^{-1} d^{-1}$ | Production rate |
| | $\rho$ | $\geq 0$ | $n_R d^{-1}$ | Reversion rate |
| (H5)<br>Reduced<br>viral<br>replication | $\pi$ | $[0,1]$ | - | Sensitivity to interferon |
| (H7, H6)<br>Natural<br>killer cell<br>dynamics | $\nu_N$ | $\geq 0$ | $n_K n_V^{-1} d^{-1}$ | Recruitment rate |
| | $\delta_N$ | $\geq 0$ | $d^{-1}$ | Clearance rate |
| | $\psi$ | $\geq 0$ | $n_I n_N^{-1} d^{-1}$ | Attack rate |
| | $\zeta$ | $\geq 0$ | $n_I n_F^{-1} d^{-1}$ | Attack rate (implicit) |

**Table 3.**
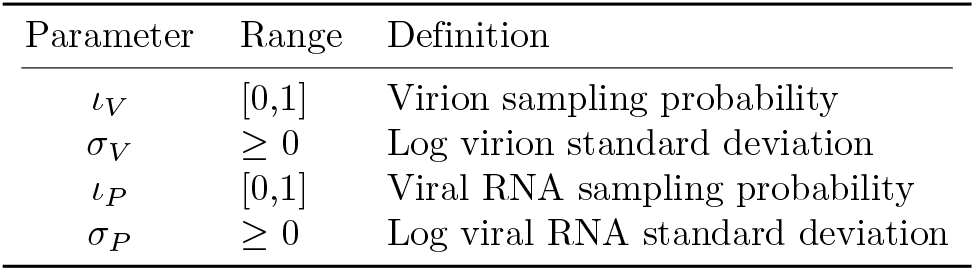
Model parameters with definition for the joint observation model for viral titers and RNA.

Viral infection data (titers and RNA) are right censored to account for the finite limit of the detection associated with viral quantification assays. Further, these detection thresholds (*DT*) differed between virus type (H3N2 and SARS-CoV-2) and data type (viable virus, RNA; Fig. 2). We account for right censoring in the data by incorporating a threshold in the observation model. If the sampled trajectories drop below the threshold *DT*, according to Eq. 40, the likelihood is evaluated at the data-type specific threshold *DT*_*n*_. If not, the likelihood is evaluated at the sampled values, *Y*_*n*_(*t*, Θ).

**Fig 2.**
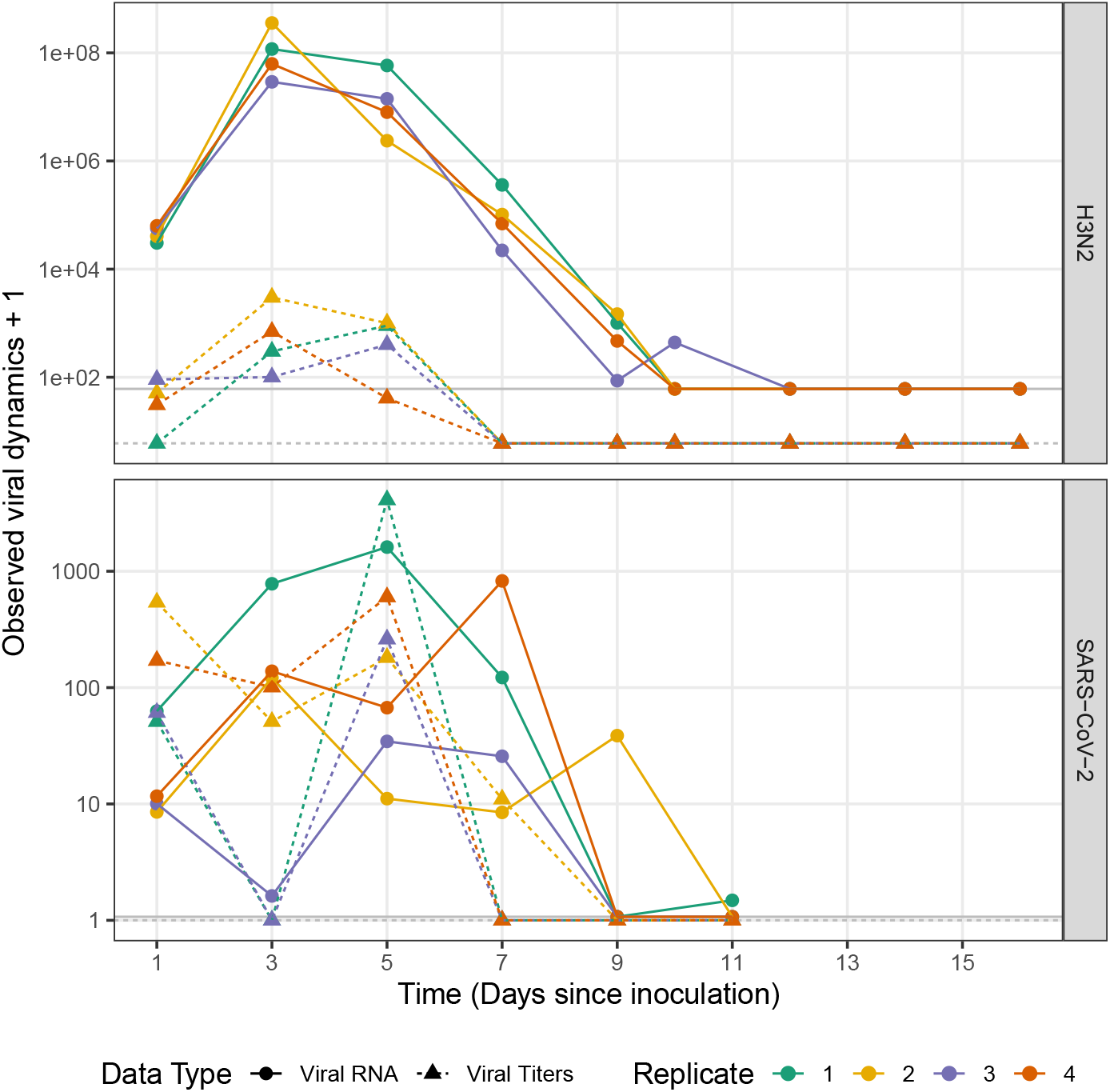
Experimental data from ferret infection experiments. Facets represent within-host viral dynamics of H3N2 influenza (top) and SARS-CoV-2 (bottom) collected from nasal washes of 4 individual ferrets (colors). Viral RNA data is represented using points and solid lines; Viral titer data is represented using triangles and dotdashed lines. Different detection threshold (dt) against virus-data type combinations are represented using solid and dotdashed lines for viral RNA and titer respectively. X-axis represents time in days since ferrets were inoculated.

#### Numerical optimization

Maximization of the likelihood (Eq. 40), is carried out using a differential evolution, stochastic optimizer routine offered by R package *DEoptim* [48]. For all hypotheses, a large population of parametric configurations (g = 500) is used to initialize the optimizer. This population is generated using a Sobol design to ensure efficient sampling of the parametric space [43, 44]. Trial mutants *v*_*i,g*_ under this computational strategy are generated using three randomly chosen candidates from the original population such that *v*_*i,g*_ = *x*_0,*g*_ + *F* (*x*_1,*g*_ − *x*_2,*g*_) [49]. We assume a high cross-over probability (0.9) and a moderate scaling factor (0.6) [50]. Owing to the complexity of the estimation problem, a very strict convergence criterion is set to ensure convergence to a global maximum (steptol = 1500 iterations, reltol = 10^−8^).

#### Quantifying goodness of fit

We use Akaike information criterion corrected for a small sample (*AICc, n* = 8 and 6 for IAV H3N2 and SARS-CoV-2, respectively) to assess the relative goodness of fit among the 7 competing models:

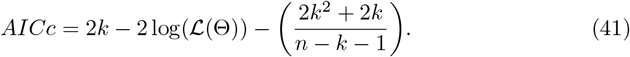

Here, ℒ (Θ) is the likelihood function from Eq. 40 and *k* denotes the number of model-specific parameters. At a 5% level of significance, an *AICc* value of 2 was taken to signify a statistical difference in the comparative agreement among competing models. Δ*AICc* = 0 was taken to denote the best mechanistic explanation of the observed dynamics.

#### Quantifying uncertainty in parameter estimates

The method of parametric bootstrap is used to estimate uncertainty in parameter MLEs. For this procedure 100 synthetic trajectories are simulated for each virus-model combination using the MLE. A global search optimizer, with less stringent convergence criterion (steptol = 500 iterations, reltol = 10^−8^), is used to re-estimate model parameters for every synthetic time series. This generates a bootstrapped distribution of parameter estimates surrounding the MLE. 95% confidence bounds for parameter uncertainty are ascertained by calculating the 2.5^th^ and 97.5^*th*^ percentiles of the bootstrapped distribution of model parameters.

### Effect of antivirals

#### Antiviral dynamics

Antivirals are chemical or biological agents that impede the within host transmission process. They may achieve this via different mechanisms: (I) rendering naïve target cells partially resistant to infection; (II) inhibiting viral replication, or blocking viral fusion, or promoting virus neutralization, thereby reducing, on average, the viral output from productively infected cells; and (III) increasing the cytotoxicity associated with infected cells by assisting in the identification and flagging for resident antagonistic cells like macrophages and natural killer cells. To study the differential effect of these qualitatively distinct antiviral mechanisms, we construct a model incorporating an antiviral exhibiting first-order kinetics as shown in Eq. 42-49.

After the initiation of therapy at time *t*_*dose*_, antiviral molecules infiltrate both susceptible target cells (*T*) and productively infected cells (*I*) at the rate *β*_*A*_ thus giving rise to uninfected target cells (*T*_*A*_) and productively infected cells (*I*_*A*_) in an antiviral phase. This allows for an explicit description of the effects of the antivirals on the transmission process. We allow for periodic administration of the antiviral at *t*_*dose*_ intervals.

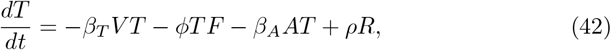

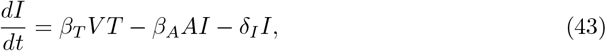

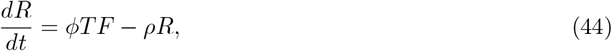

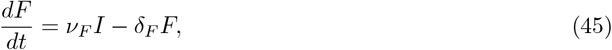

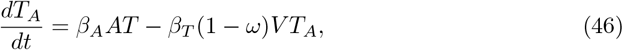

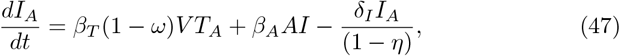

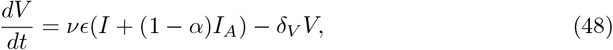

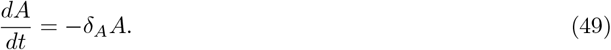

Here *δ*(*t, t*_*dose*_) is the Kronecker delta and *x* is an integer. Definitions on antiviral model compartments and parameters are provided in tables 4 & 5.

**Table 4.** Model compartments for the model of antiviral action.

| Compartment | Definition |
| --- | --- |
| $T$ | Target epithelial cells, invasion sites |
| $I$ | Infectious epithelial cells, replication sites |
| $V$ | Viral titer |
| $R$ | Refractory epithelial cells |
| $F$ | Type 1 interferon ($\alpha$ and $\beta$) |
| $T_A$ | Target epithelial cells, anti-viral phase |
| $I_A$ | Infectious epithelial cells, anti-viral phase |
| $A$ | Anti-viral concentration |

As described above, we consider three separate mono-therapies that represent qualitatively distinct effects on *T*_*A*_ and *I*_*A*_. First, antiviral target cells (*T*_*A*_) have reduced susceptibility to infection, denoted by (1 − *ω*; see Eq. 46). Second, the *per capita* baseline infectious period of *I*_*A*_ cells (1*/δ*_*I*_) is reduced by a factor of (1 − *η*), thereby increasing the average cytotoxicity of antiviral-containing infected cells (see Eq. 47). Third, the average viral production per *I*_*A*_ cell is reduced by a factor (1 − *α*) thereby reducing the viral burst size per productive cycle (see Eq. 48).

#### Antiviral performance indicator

Measure of therapeutic performance is ascertained using the virological efficacy index, which we define as the percent reduction in the area under the viral curve under treatment (*AUV C*) compared to the the area under the viral curve for the control (*AUV C*_*c*_) [51]. Area under the curve in both cases was approximated using the trapezoidal rule implemented in the R package AUC [52].

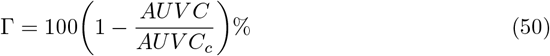

## Results

### Viral dynamics

There are virus-specific differences in the quantified viral dynamics across individual animal replicates for H3N2 and SARS-CoV-2 (Fig. 2). H3N2 shows a more rapid viral take off that is consistent among individual animals, reaching an earlier peak (day 3) relative to SARS-CoV-2 that exhibits greater inter-individual variability in initial dynamics and reaches a maximum viral load typically on day 5. Both viruses fall below the detection threshold by day 11. Unlike H3N2, we observe a multi-modal infection curve for SARS-CoV-2. We examine whether sex-specific differences account for this variation in viral kinetics but find no discernible pattern (Fig. S.1).

### Parameter sensitivity to model outcomes

To explore how simulated dynamics respond to changes in model parameters, we conduct a systematic sensitivity analysis calculating partial rank correlation coefficients (PRCC) between combinations of parameter and three model outcomes. We find that nearly all of the PRCC values were significant (*p* − *value <* 0.05). However, many of the model parameters have small effect sizes (*PRCC <* 0.1). Among the model parameters, Initial viral load (*V*_0_) and the basic reproductive number (*R*_0_) have the largest *PRCC* associated with viral decay rate and the peak viral load (see Fig.S.2). It should be noted that *R*_0_ is a composite derived quantity and the effects of individual parameters (see Eq.39) are not considered individually mainly to assess the system sensitivity while reducing the dimension of the parameter space. We do not observe large effect sizes for immunity parameters (*PRCC <* 0.2, Fig.S.2) in generating the chosen model outcomes. One reason leading to this result could be attributed to the fact that simple model outcomes, principally the initial peak size, timing and decay slope, are short-term phenomena that are driven by the processes associated with transmission and not immunity. Immune mechanisms tend to generate more complex dynamics on a longer time scale.

### Statistical inference with transmission models

We use the viral data obtained by both PFU/FFU and qRT-PCR assays (Fig. 2) in likelihood-based statistical inference to evaluate the evidence in support of each hypothesis (as represented in Eq. 1–35). As depicted in Fig. 3, all of proposed models can provide a good explanation of the data. However, as indicated by ΔAICc values, we find individual models for H3N2 and SARS-CoV-2 that best fit the data (Fig. 3). We further quantify model-data agreement for the best-fitting model for each virus and data type: H3N2 (*R*^2^ for viral RNA = 99.81%, viral titers = 85.22%; Fig. S.3) and SARS-CoV-2 (*R*^2^ for viral RNA = 92.68%, viral titers = 49.20%; Fig. S.3).

**Fig 3.**
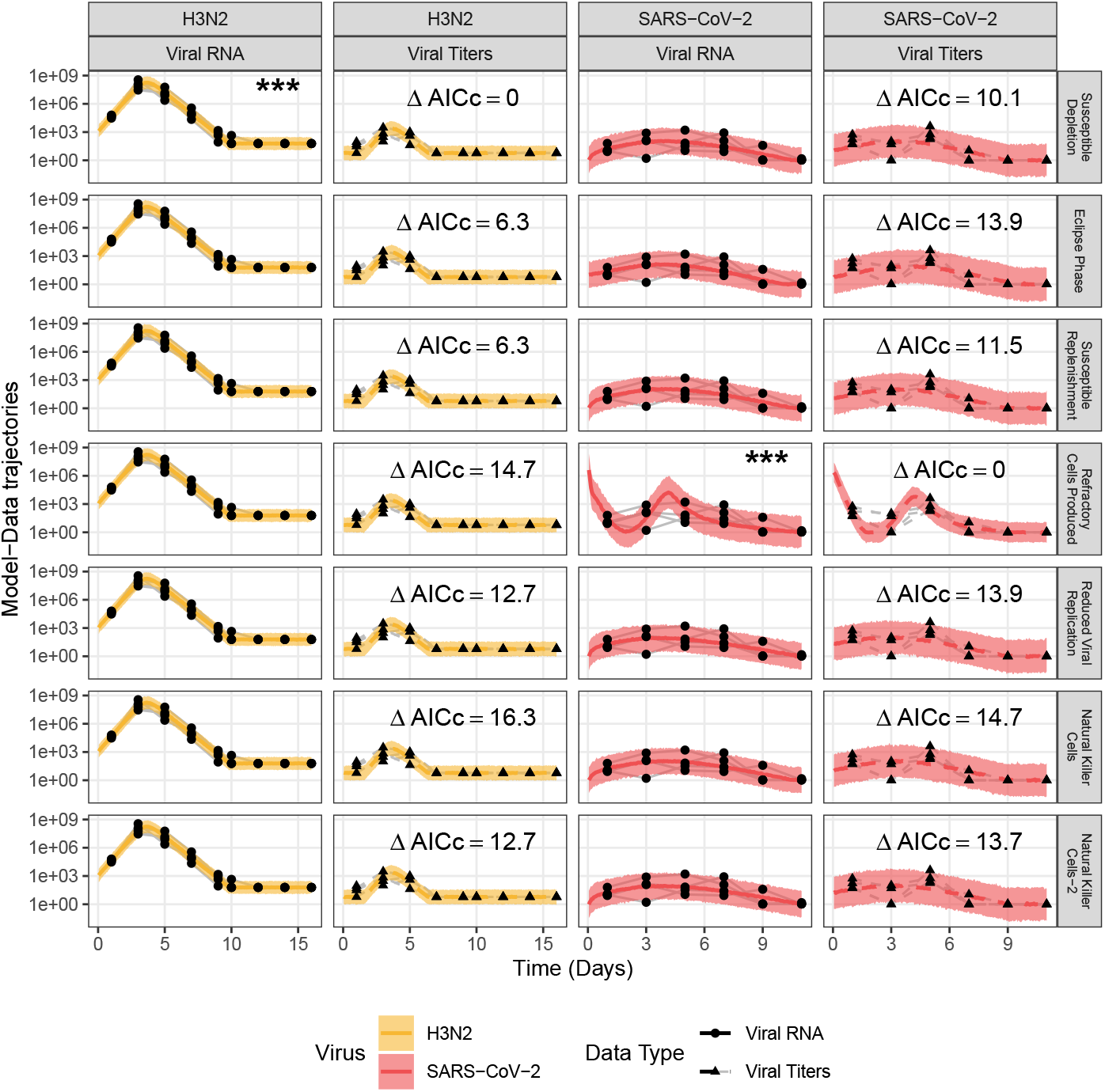
Relative fits of within-host models to virus dynamics of H3N2 and SARS-CoV-2. Models trajectories were fit to time series of virus titer (circles-solid lines) and RNA (triangles-dashed lines). Lines represent simulated median of the log-normal probability model and ribbons represent the 95% prediction intervals for H3N2 (golden) and SARS-CoV-2 (red) respectively. (***) denotes the best fitting model denotes the best fitting model to the observed dynamics (Δ*AICc* = 0)

The supported hypothesis for the two viruses is different. In the case of H3N2, the most parsimonious explanation is the simplest model (“susceptible cell depletion”) which is sufficient to explain the observed viral dynamics (ΔAICc = 0). This model has been previously documented to capture the single viral peak that is characteristic of H3N2 infections [53]. More surprising however is that all other models, initialized over a wide range of parameter hyperspace, essentially converged to the best-fitting model. Tables 6 and S.1 present maximum likelihood parameter estimates (MLEs) and their corresponding confidence intervals for each hypothesis. Relative differences in AICc values of the competing models for H3N2 infection are ascribed to the penalties incurred on account of higher parametric degrees of freedom with the more complex dynamical explanations. We estimate a modest degree of intercellular transmissibilty for H3N2 (*R*_0_ = 1.64 (1.27, 1.98)), a high probability of sampling and low variability in both viral RNA (*ι*_*P*_ = 1.00 (0.01, 1), *σ*_*P*_ = 0.79 (0.6, 0.94)) and viral titer (*ι*_*V*_ = 0.66 (0.26, 0.98), *σ*_*V*_ = 0.90 (0.68, 1.09)).

**Table 5.**
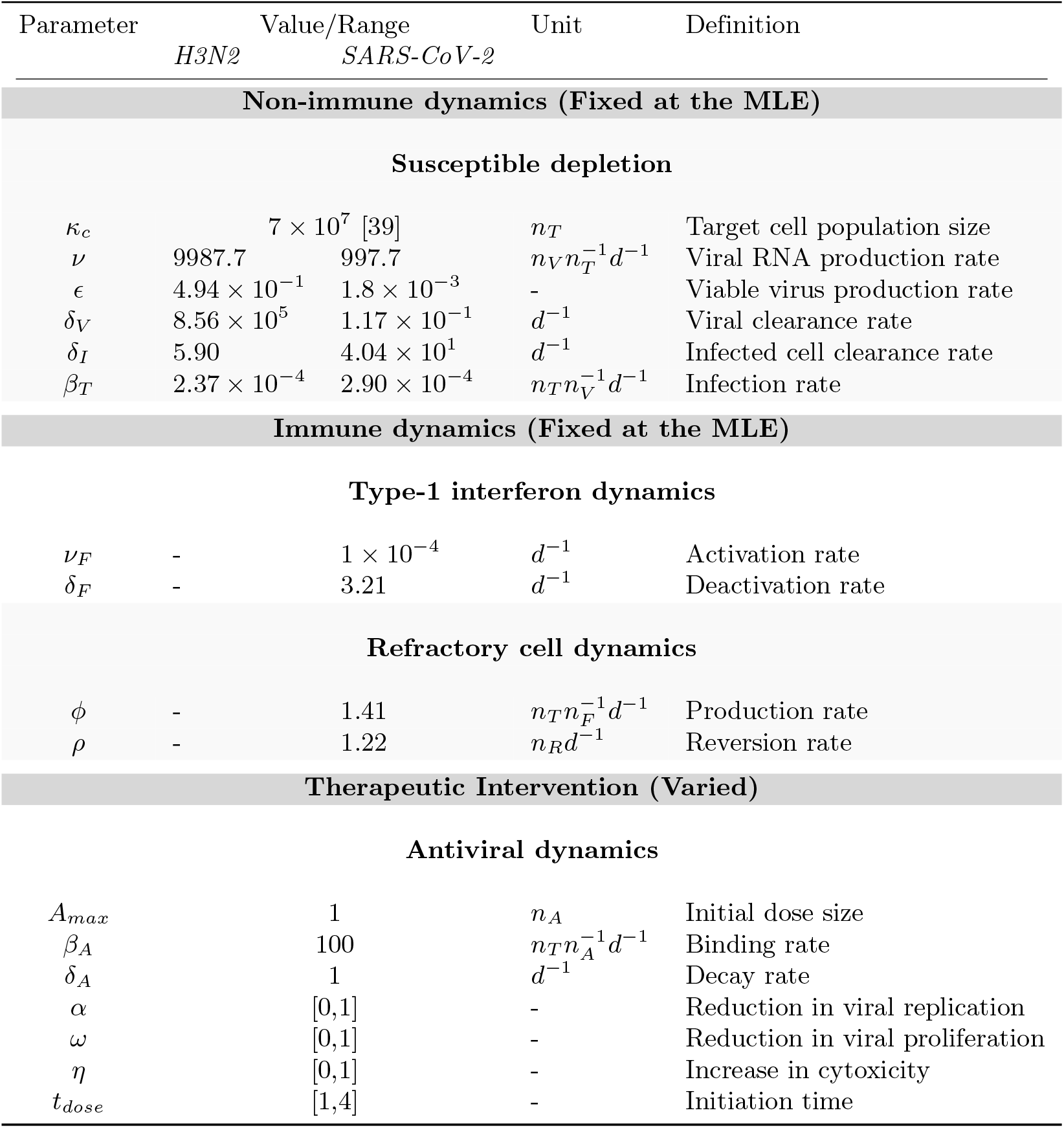
Model parameters with definition and units for the full model of infection and immunity. Parameters are grouped by the hypotheses tested in this article and arranged in the increasing order of biological complexity. Individual model formulations are specified in Eq. 42-49

**Table 6.**
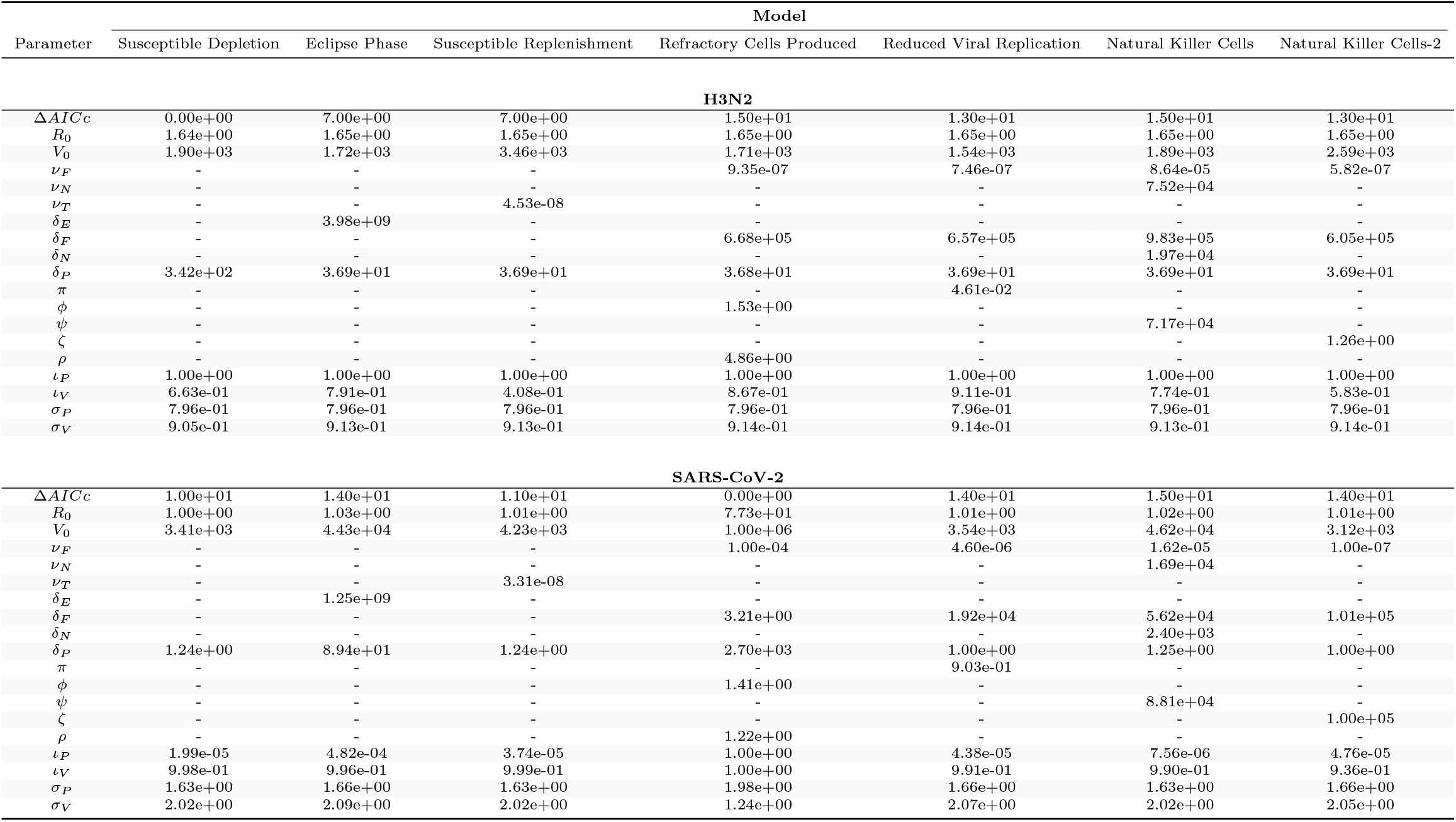
Maximum likelihood estimates (MLE). Model specific parameter MLE’s were obtained by maximizing the likelihood function.

Relative to H3N2, the analysis of SARS-CoV-2 data supports the “increased production of refractory cells” hypothesis as the most parsimonious explanation (ΔAICc = 0). We estimate a nearly 50-fold higher degree of intercellular transmissibility (*R*_0_ = 77.30 (22, 163)), compared to H3N2. Refractory cells are estimated to be produced at a rate of 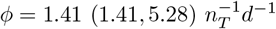 but also a rapid reversion rate such that the refractory phase lasts for 0.82 (0.72, 1.37) 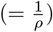 days. Similar to H3N2, we observe a low degree of sampling bias for both data types (*ι*_*P*_ = 1 (0.12, 1), *ι*_*V*_ = 1 (0.4, 1)), however, a much higher degree of variability (*σ*_*P*_ = 1.98 (1.47, 2.51), *σ*_*V*_ = 1.24 (1.06, 1.64)) is required to explain the variance in the observed infection dynamics. This, we propose might be a consequence of the high degree of individual-level variation exhibited by SARS-CoV-2 replicates unlike H3N2 (Fig. 3).

### Therapeutic outcomes under differential antiviral interventions

The best-fitting models for each virus are used to assess the effects of different antiviral therapies (see Eq. (42)-(49)) on viral control and treatment. The model is fixed at a moderate value of antiviral reactivity 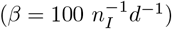 and duration (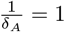1 day) and we assume a daily pulsed administration of an antiviral that enhances cytotoxicity by killing the infected cells under antiviral influence(Fig. 4). The effectiveness of model outcomes is ascertained using Eq. 50. We then qualitatively demonstrate the relationship between the timing of the initiation of therapy and the efficacy of the antiviral, defined as the increase in the cellular toxicity (*η*). We observe that a smaller value of *η* is equally effective provided the intervention is initiated in the early phase of the infection. Owing to the higher values of within-host transmissiblilty (*R*_0_ = 77.3) and cytotoxicty of cells infected with SARS-CoV-2 (*δ*_*I*_ = 44 *d*^−1^), we observe that generally a more efficacious antiviral (*η* ≥ 0.5) is required to produce an effect comparable in the therapeutic outcome in treatment of H3N2 (*R*_0_ = 1.64, *δ*_*I*_ = 4 *d*^−1^).

**Fig 4.**
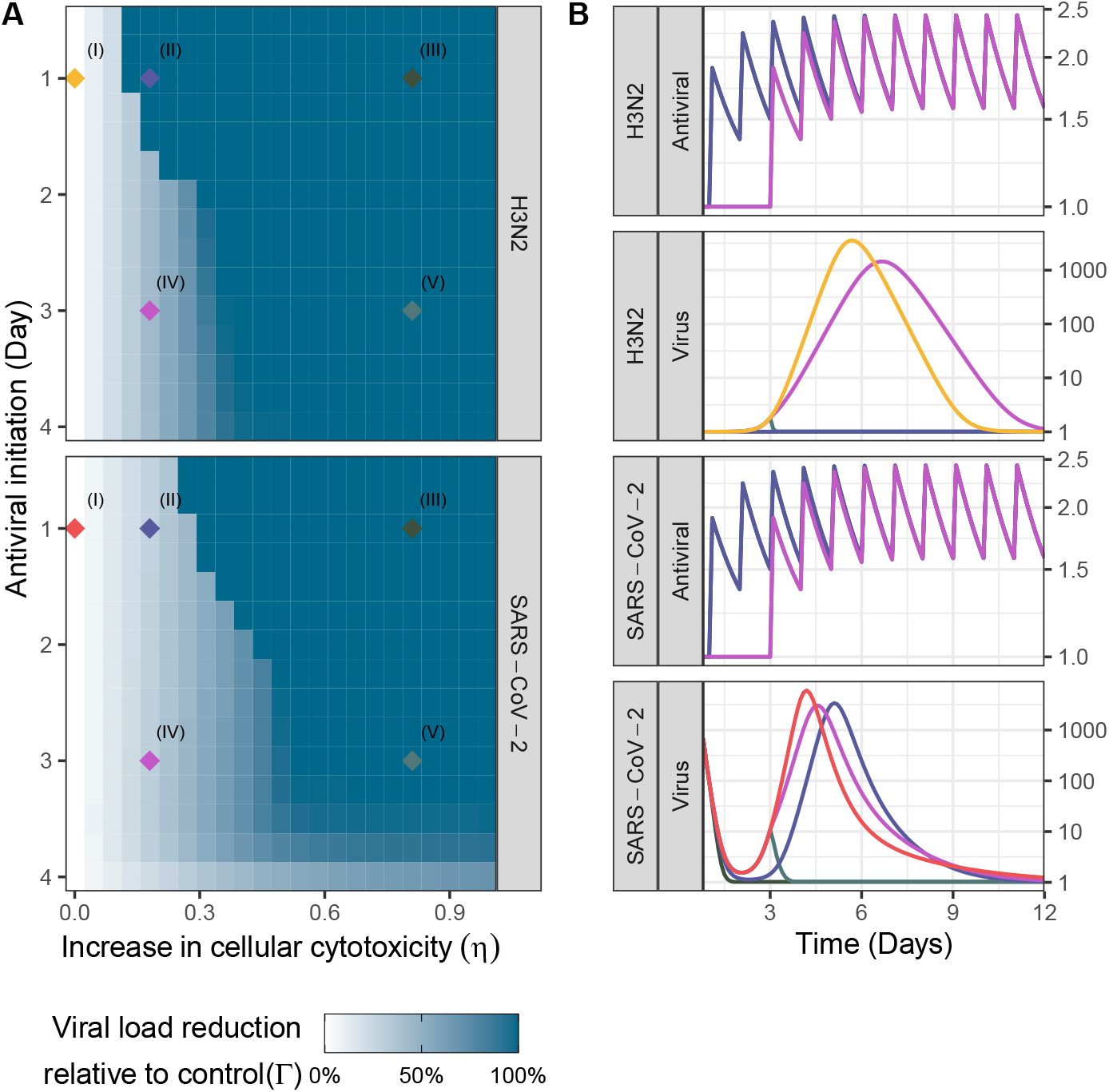
Relationship between effectiveness and initiation time for antiviral enhancing cellular toxicity on the viral dynamics of H3N2 and SARS-CoV-2. (A) Effectiveness of the antiviral therapy expressed as the reduction in of the area under the viral curve relative to the control region. (B) Simulated viral and anti-viral dynamics for regions (I-V) in (A). Control trajectories (region I, *η* = 0) of virus dynamics of H3N2 (golden) and SARS-CoV-2 (red) were generated using the parameter MLEs of best fitting models. For treatment (regions II-V), we assume a daily pulse of unit antiviral drug (*Amax* = 1) with moderate antiviral binding rate 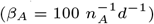 and duration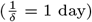.

Surprisingly, when comparing between modes of antiviral action (as represented by the parameters *ω, η* and *α*), we find little noticeable difference in efficacy (Fig. S.4). To explore this, we analytically derive an antiviral impact factor using the next generation operator [42]. For the full antiviral model described in Eq.42-49, we derive the within-host effective reproductive number (*R*_*A*_, for details on derivation see Appendix A1),

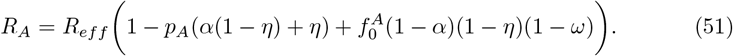

Here, *R*_*eff*_ = *R*_0_*f*_0_ is the effective reproductive number at the time of the antiviral administration. 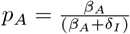, is the probability of getting bound by the antiviral; 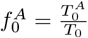, is the fraction of target cells bound by the antiviral 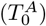 to the susceptible target cells (T_0_). This expression (Eq. 51) calculates the average number of infected cells produced by a single infected cell in the instant following administration of the antiviral.

We define antiviral impact (*ξ*) as the reduction in the within host transmissibilty of the virus in the presence of an antiviral (*R*_*A*_) as compared to a baseline (*R*_*eff*_). Then from Eq. 51 we get:

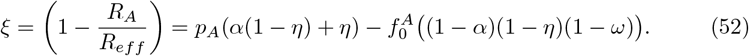

From this equation, we observe that the antiviral impact depends not only on the efficacy of the drug *i*.*e*. the absolute values of *α, η*, and *ω*, but also is a product of both *p*_*A*_ (within-host reactivity and duration of the antiviral) and 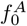 (a proxy for the dependence of antiviral impact on the timing of the antiviral dose). For this simple case, we find that the effect of the modes of antiviral action appear multiplicatively and cannot be separated. This, we submit, is the main reason for the lack of qualitative differences among the modes of antiviral action reported in Fig.S.4.

## Discussion

In this study, we explore the potential role of different mechanisms in explaining the observed within-host viral dynamics of influenza A H3N2 and SARS-CoV-2 infections in ferrets. By formulating a suite of hypotheses expressed as mathematical models, we use likelihood-based inference to arbitrate among these competing mechanisms. We find that the viral kinetics of H3N2 are most parsimoniously explained by the simplest model assuming only susceptible depletion. In contrast, the SARS-CoV-2 data support a more complex explanation involving the generation of refractory susceptible target cells stimulated by type I and III interferons. Further, we find that SARS-CoV-2 has a much higher rate of within-host proliferation (as summarized by *R*_0_) than H3N2. This striking difference may, we propose, be a potential explanation for the different conclusion reached regarding the putative association with the host immune system. Given the inconsistent and stuttering initial phase of the viral curve for SARS-CoV-2 (Figs. 2 & 3), the model’s preferred explanation is an initially large population of refractory cells that eventually reactivate susceptibility as the cytokine production declines due to reduced infectious cells. This mechanism introduces a brief delay in the dynamics with the infection of these newly susceptible target cells eventually contributing to viral proliferation. Previously, a similar finding was reported to explain the bi-modality and the prolonged plateau in peaks in observed equine influenza A (H3N8) [54] and human influenza A (H1N1) dynamics [55].

In contrast, the lower *R*_0_ value for influenza stimulates a weaker interferon response. Interestingly, a lower proliferation rate for H3N2 thus facilitates rapid viral spread with termination of infection brought about by susceptible depletion. This conclusion is consistent with studies of viral dynamics at during the early stages of an infection in humans [56, 57]. We note, however, that our conclusion does not preclude the involvement of the host immune system in modulating H3N2 infection. We speculate that the dynamical simplicity of these data may affect the potential for identification of more complex mechanisms. Resolution of this issue will require the incorporation of additional parallel immunological signals in such analyses [58, 59].

Plaque-based assays (PBA) generate statistically accurate estimates of cellular infection burden by quantifying viral titers through the number of viable virions [60, 61]. However, with the manual assessment of plaque-forming units (PFU) often associated with PBA, data generated through these methods may be noisy. A statistically precise measure of within-host viral burden, on the other hand, can be obtained using sampling methods based on quantitative polymerase chain reaction (qPCR) [62]. These methods generate data that can be orders of magnitude higher than the viable virus titer (*eg*., For foot-and-mouth disease, the PBA:qPCR ratio is ∼ 1 : 1000 [63]). Such discrepancies in the estimated viral titer can lead to an over-estimate of the proliferation potential of the virus or make comparison of different studies challenging. For this reason, our models simultaneously account for both qPCR and viral titer data.

To examine the potential control of these viruses with antivirals, we formulate models that permitted different modes of antiviral action. Antivirals may serve to: reduce susceptibility of target cells, lower viral production by infectious cells and increase infectious cell mortality. We find no qualitative difference among the three modes of action (see Fig.S.4). A similar result has been discussed in while devising antiviral therapies for closely related coronaviruses [2]. Our theoretical analysis explained how antiviral effectiveness is determined by a product of the three individual antiviral modes of action. Hence, their mono-therapeutic effects are qualitatively indistinguishable, as we show in Eq. 52.

In this study we have attempted to understand the within-host interactions between invading viral pathogens and the host’s innate immune system. Further, we qualitatively describe the differential action of antiviral therapies by leveraging on the pre-existing immune interaction for H3N2 and SARS-CoV-2 infections.

## Acknowledgments

This project has been funded in part with Federal funds from the National Institute of Allergy and Infectious Diseases, National Institutes of Health, Department of Health and Human Services, under Contract No. HHSN272201400004C (NIAID Centers of Excellence for Influenza Research and Surveillance, CEIRS), Contract No. 75N93021C00018 (NIAID Centers of Excellence for Influenza Research and Response, CEIRR) and from the National Science Foundation under Grant No. DGE-1545433 (IDEAS training program). We would like to thank Leila Beheshtivayeghan for her contribution to the schematic representation of the infection model.

## Supporting information

## Appendix A1

This appendix documents the derivation for the antiviral effective reproductive number presented in Eq.51. We use the method of next generation operator (NGO) [42] to derive the quantity *R*_*A*_. MATLAB package *symbolic toolbox* [64] was used to simplify matrix and algebraic expressions. The NGO method applied is described in the following steps:

1. From the system of differential equations documented in Eq. 42-49, compartments that contribute to the productions of novel infected cells were identified. In this case, the relevant compartments are - (1) productively infected cells (*I*), (2) productively infected cells bound to the antiviral molecules (*I*_*A*_) and (3) free virions (*V*)
2. For each of these compartments, two rate vectors were defined –

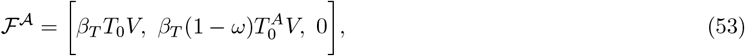

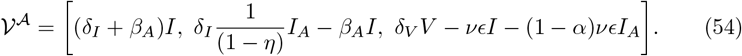

ℱ ^A^ defines the vector of rates at which new infected cells are produced. 𝒱 ^A^ defines the vector of the flow in and out of the infected cell compartments.
3. Jacobian matrices were then defined using vectors in Eq. 53-54. To linearize the system in Eq. 53-54, it was assumed that the susceptible target population and anti-viral bound target population were are at constant values *T*_0_ and 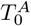 respectively i.e. the two matrices evaluated at the infection free equilibrium. **F**^*A*^ is such that every element 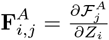, and **V** ^**A**^ is such that every element 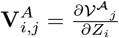, where *Z* = [*I*_*A*_, *I, V*] is the vector of relevant state variables that generate novel infections arise. We express the Jacobian matrices to be –

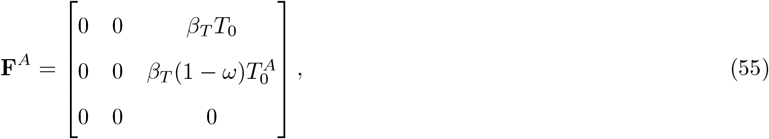

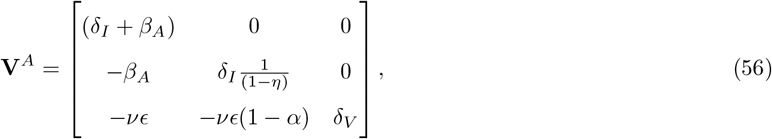

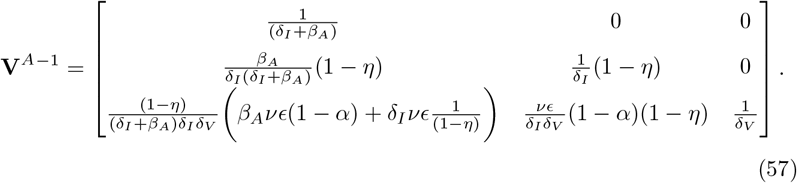 Here, **V**^*A*−1^ is the inverse of the **V**^*A*^ matrix which was used in the calculation of the next generation matrix.
4. We then proceeded to calculate the next generation operator as –

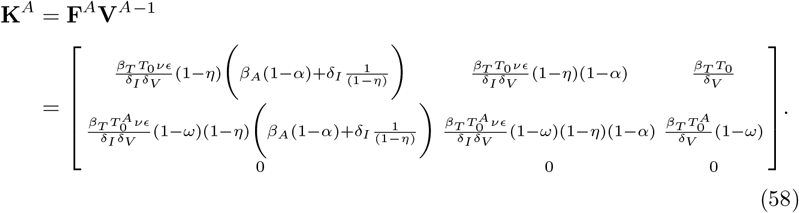
5. The largest eigen value of matrix **K**^*A*^, calculated by solving the characteristic equation, |**K**^*A*^ − Λ**I**| = 0, represents *R*_*A*_. Here, **I** is the identity matrix.

Following the NGO method and simplifying the largest cubic root of the characteristic equation,

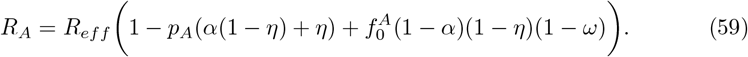

which is reported in Eq. 39

## Supporting figures

**Fig S.1.**
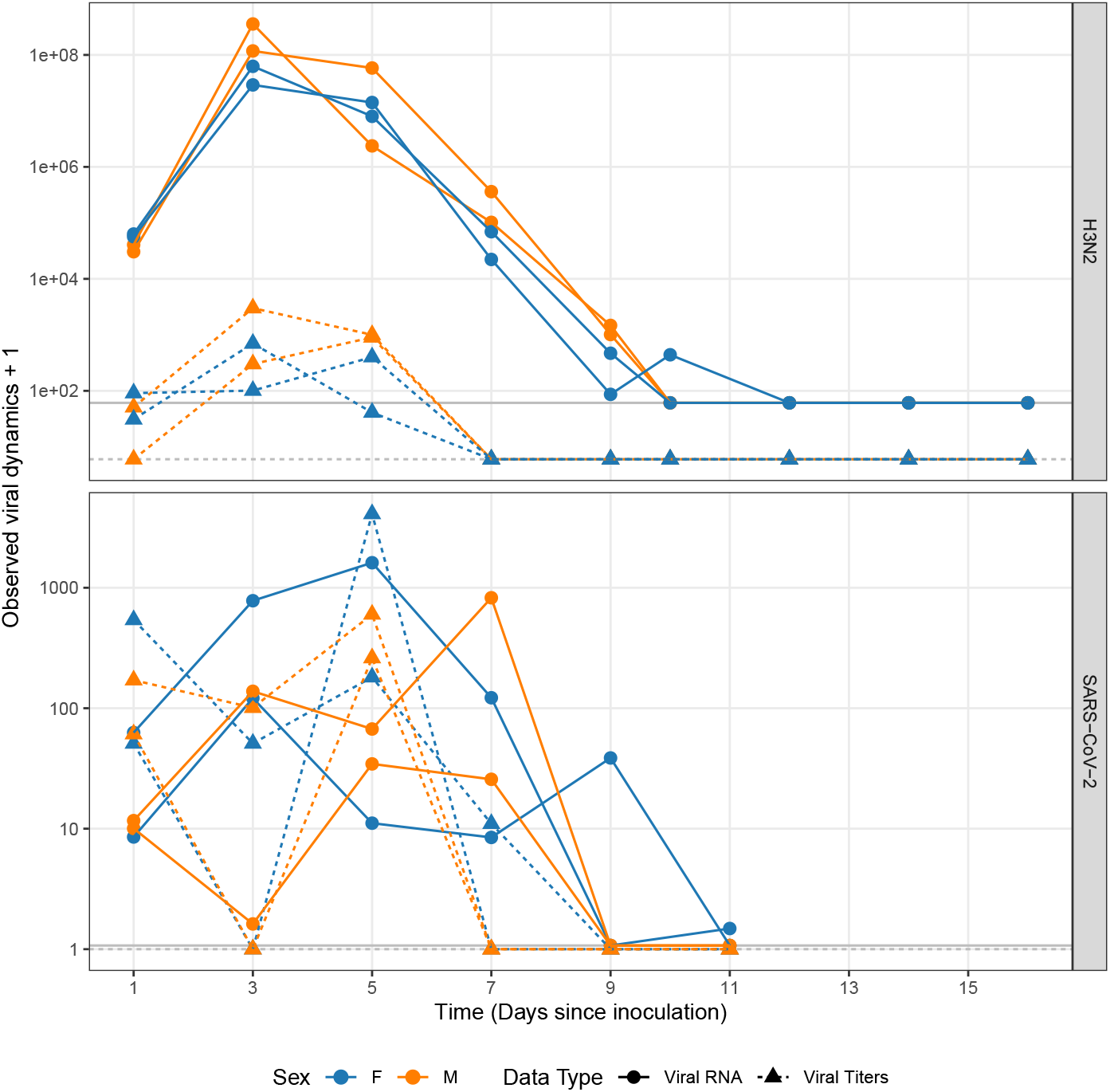
Experimental data from ferret infection experiments stratified by the sex of the animal. Facets represent within-host viral dynamics of H3N2 influenza (top) and SARS-CoV-2 (bottom). Viral RNA data is represented using points and solid lines; Viral titer data is represented using triangles and dotdashed lines. Different detection threshold (dt) against virus-data type combinations are represented using solid and dotdashed lines for viral RNA and titer respectively. X-axis represents time in days since ferrets were inoculated.

**Fig S.2.**
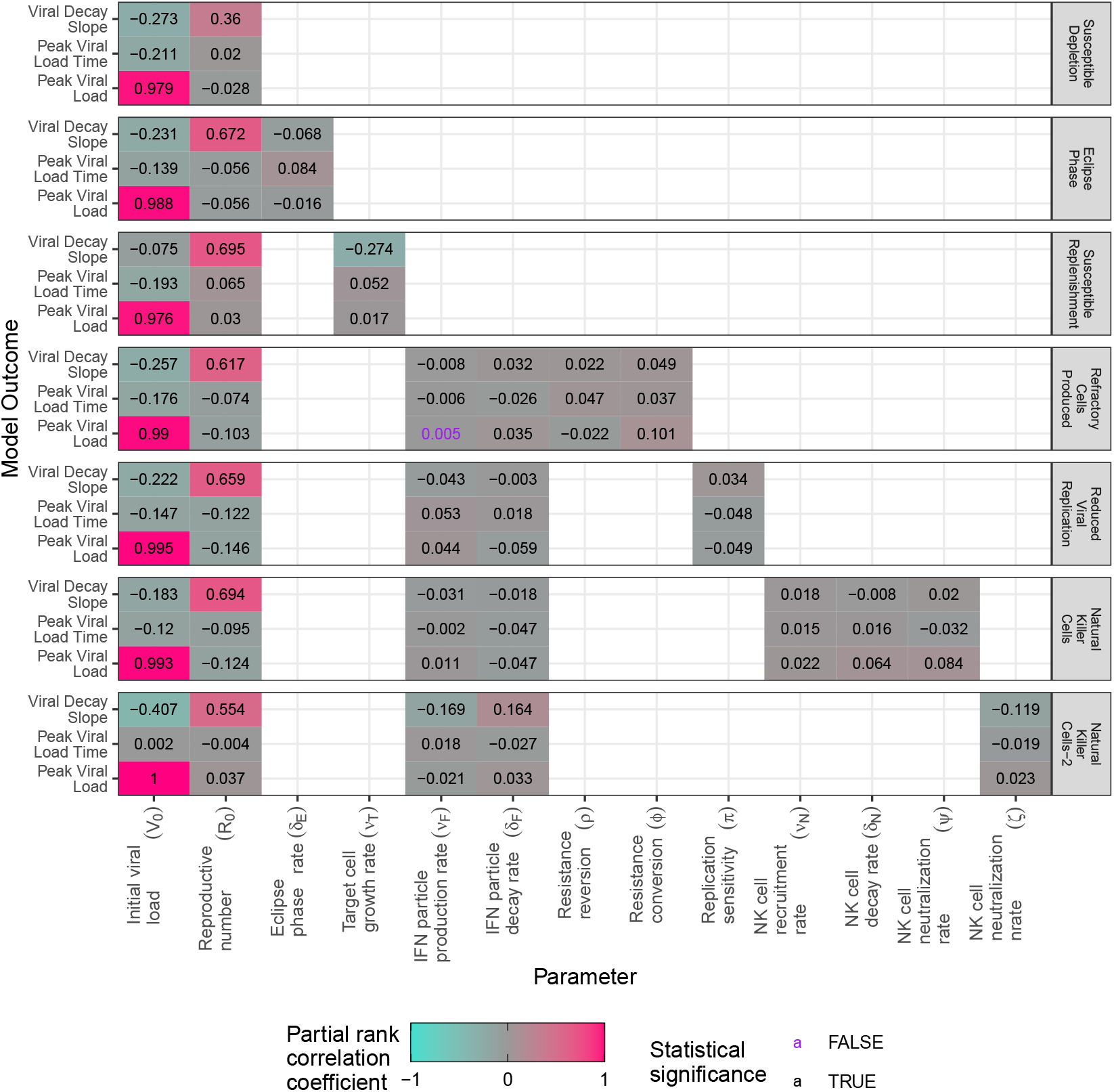
Sensitivity analysis with hypothesis specific model parameters. Partial rank correlation coefficients (PCC, color) were calculated to assess sensitivity of parameters in generating model outcomes. 5000 parameter configurations were generated using a Sobol sampling scheme. Bootstrapped confidence intervals were obtained by uniformly sampling with replacement and generating 150 samples. Statistical significance was ascertained at 5%, Bonferroni corrected, level of sigificance.

**Fig S.3.**
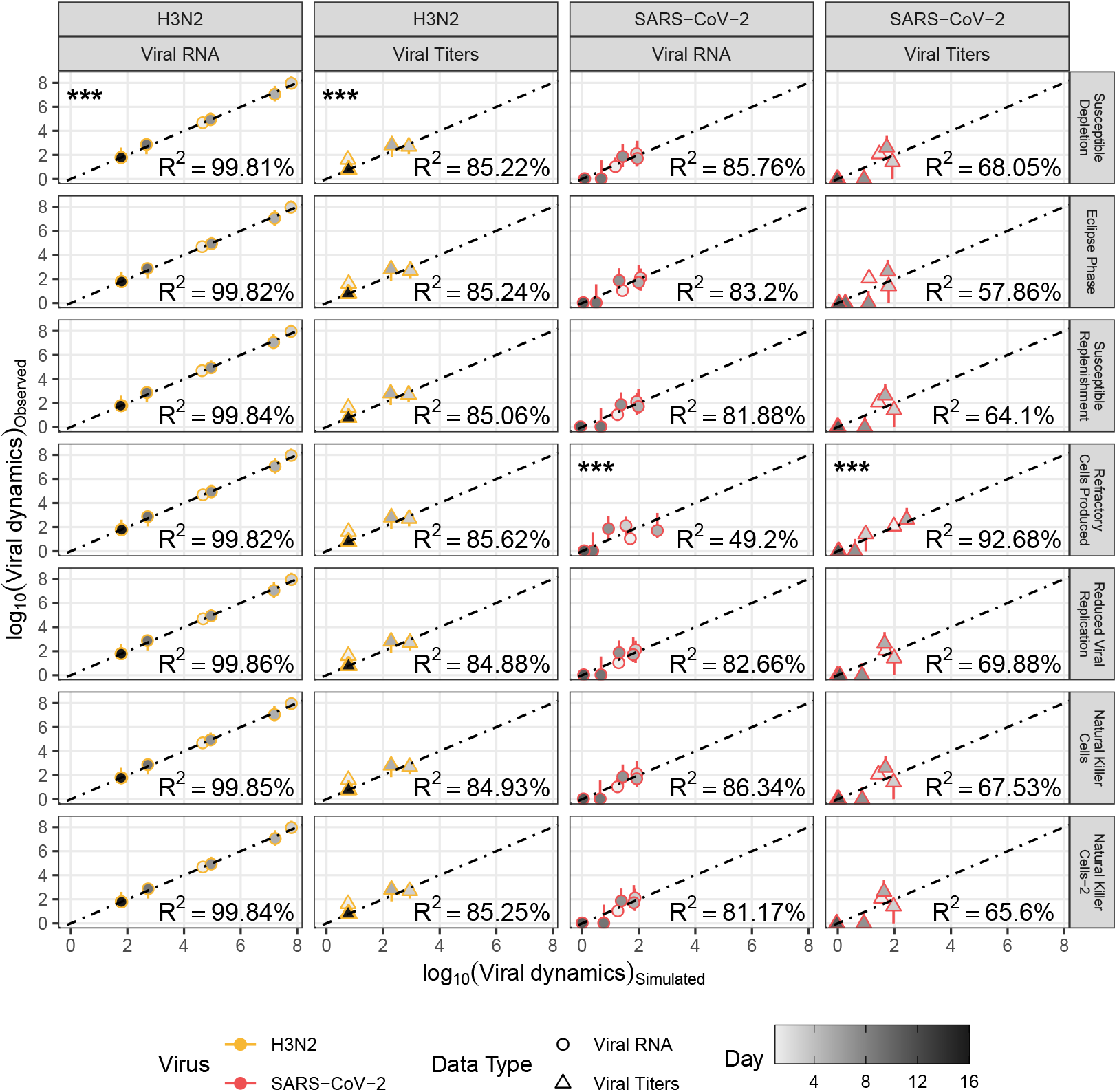
Goodness of fit for virus-data-model combination. Scatter plots show model-data agreement for viral RNA (empty circles) and viral titers (empty triangles) on a *log*^10^ scale. Error-bars represent individual level variation among replicates. Fill gradient represents time (in days) to demarcate the observed infection interval. Dotdashed line is the 1:1 reference line. For every model, coeficient of variation (*R*^2^, reported inset) was calulated for each data-type at the medium of the replicate ditribution to quantify the proportion of variation explained by the models. (***) represents the best fitting model based on the AICc comparison.

**Fig S.4.**
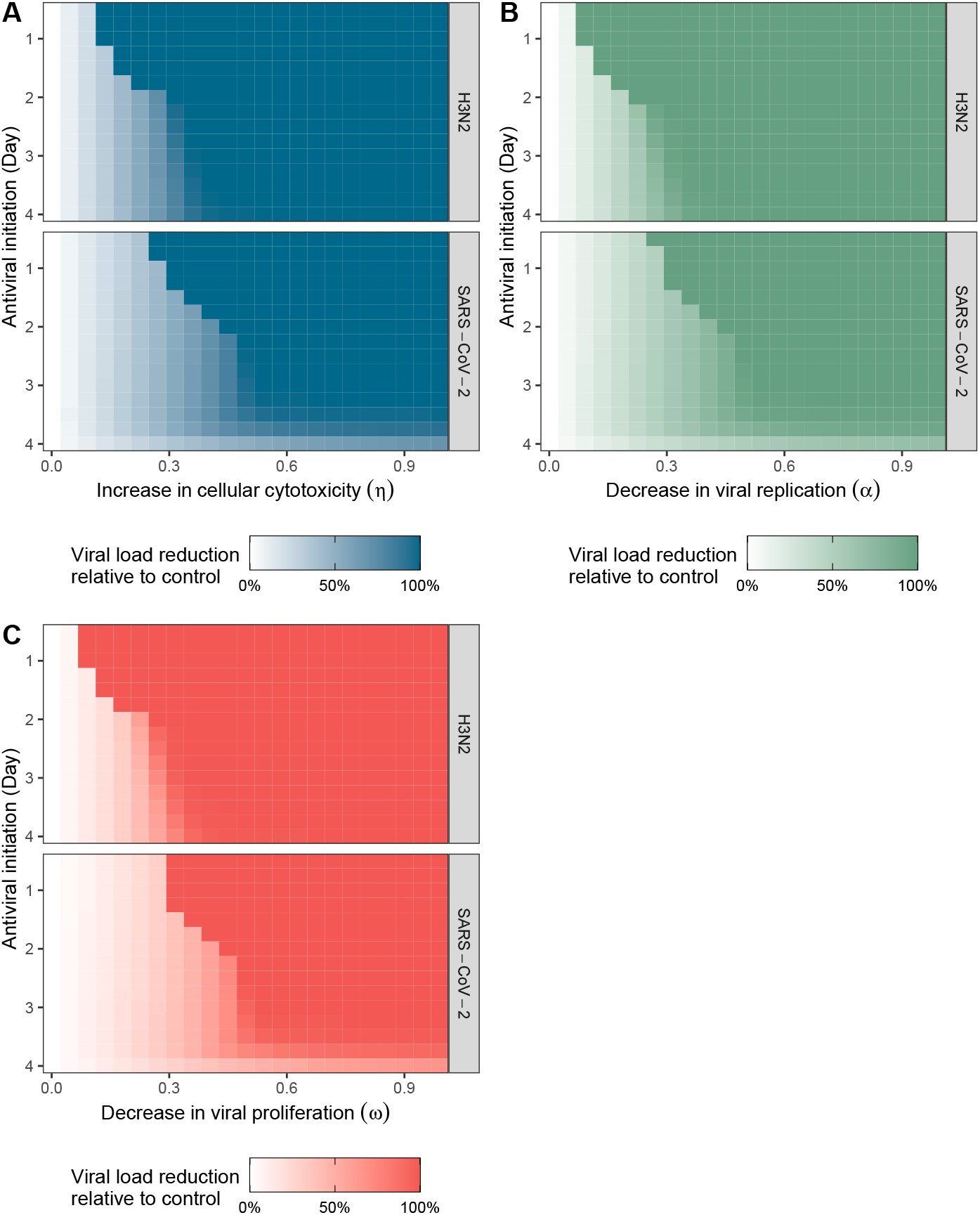
Simulation study reflecting the relative action of antiviral effectiveness, initiation time and type on the viral dynamics of H3N2 and SARS-CoV-2 with a pulse dose administrative strategy. (A, C) Effectiveness of the antiviral therapy expressed as the reduction in of the area under the viral curve relative to the control region. (B, D) Viral and anti-viral dynamics simulated for regions (1-5). Control trajectories, region (1), for the two viruses were generated using the parameter MLEs of two best fitting models and simulating dynamics for 12 days. For the remainder of regions, a moderate antiviral binding rate 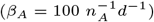 and duration 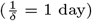 were assumed during this study.

**Table S1.** Maximum likelihood estimates (MLE) and 95% confidence intervals. Model specific parameter MLE’s were obtained by maximizing the likelihood function. Uncertainty in the MLE was ascertained using parametric bootstrap.

| Parameter | Model |  |  |  |  |  |  |
| --- | --- | --- | --- | --- | --- | --- | --- |
|  | Susceptible Depletion | Eclipse Phase | Susceptible Replenishment | Refractory Cells Produced | Reduced Viral Replication | Natural Killer Cells | Natural Killer Cells-2 |
| H3N2 |  |  |  |  |  |  |  |
| $\Delta AIC_c$ | 0.00e+00 | 7.00e+00 | 7.00e+00 | 1.50e+01 | 1.30e+01 | 1.50e+01 | 1.30e+01 |
| $R_0$ | 1.64e+00 (1.27e+00, 1.98e+00) | 1.65e+00 (1.20e+00, 1.93e+00) | 1.65e+00 (1.28e+00, 1.87e+00) | 1.65e+00 (1.22e+00, 2.05e+00) | 1.65e+00 (1.19e+00, 1.97e+00) | 1.65e+00 (9.54e-04, 1.65e+00) | 1.65e+00 (1.26e+00, 1.96e+00) |
| $V_0$ | 1.90e+03 (7.91e+02, 1.03e+04) | 1.72e+03 (5.06e+02, 7.11e+03) | 3.46e+03 (7.69e+02, 7.48e+03) | 1.71e+03 (1.86e+02, 5.66e+03) | 1.54e+03 (4.96e+02, 5.99e+03) | 1.89e+03 (1.89e+03, 3.92e+05) | 2.59e+03 (9.45e+02, 5.58e+03) |
| $\nu_F$ | - | - | - | 9.35e-07 (2.22e-07, 9.28e-05) | 7.46e-07 (2.78e-07, 9.67e-05) | 8.64e-05 (3.53e-05, 8.64e-05) | 5.82e-07 (1.82e-07, 6.54e-03) |
| $\nu_N$ | - | - | - | - | - | 7.52e+04 (7.40e+04, 7.52e+04) | - |
| $\nu_T$ | - | - | 4.53e-08 (4.53e-08, 4.34e-02) | - | - | - | - |
| $\delta_E$ | - | 3.98e+09 (2.45e+08, 9.80e+09) | - | - | - | - | - |
| $\delta_F$ | - | - | - | 6.68e+05 (1.07e+00, 9.72e+05) | 6.57e+05 (5.24e+03, 9.82e+05) | 9.83e+05 (5.73e+05, 9.83e+05) | 6.05e+05 (2.67e+05, 9.61e+05) |
| $\delta_N$ | - | - | - | - | - | 1.97e+04 (1.80e+04, 1.97e+04) | - |
| $\delta_P$ | 3.42e+02 (3.30e+00, 6.28e+02) | 3.69e+01 (3.33e+00, 7.33e+01) | 3.69e+01 (4.09e+00, 5.21e+01) | 3.68e+01 (3.52e+00, 5.71e+01) | 3.69e+01 (3.53e+00, 6.84e+01) | 3.69e+01 (3.69e+01, 5.44e+05) | 3.69e+01 (3.38e+00, 6.59e+01) |
| $\pi$ | - | - | - | - | 4.61e-02 (8.88e-03, 9.77e-01) | - | - |
| $\phi$ | - | - | - | 1.53e+00 (1.15e-01, 9.40e+01) | - | - | - |
| $\psi$ | - | - | - | - | - | 7.17e+04 (7.17e+04, 7.35e+04) | - |
| $\zeta$ | - | - | - | - | - | - | 1.26e+00 (2.20e-01, 1.33e+03) |
| $\rho$ | - | - | - | 4.86e+00 (8.10e-02, 4.70e+01) | - | - | - |
| $\iota_P$ | 1.00e+00 (1.31e-02, 1.00e+00) | 1.00e+00 (1.74e-01, 1.00e+00) | 1.00e+00 (2.08e-01, 1.00e+00) | 1.00e+00 (1.00e-01, 1.00e+00) | 1.00e+00 (6.97e-02, 1.00e+00) | 1.00e+00 (1.68e-01, 1.00e+00) | 1.00e+00 (1.66e-01, 1.00e+00) |
| $\iota_V$ | 6.63e-01 (2.58e-01, 9.77e-01) | 7.91e-01 (4.40e-01, 9.94e-01) | 4.08e-01 (3.99e-01, 9.79e-01) | 8.67e-01 (3.77e-01, 9.92e-01) | 9.11e-01 (5.09e-01, 9.91e-01) | 7.74e-01 (3.65e-02, 7.74e-01) | 5.83e-01 (3.02e-01, 9.92e-01) |
| $\sigma_P$ | 7.96e-01 (6.00e-01, 9.49e-01) | 7.96e-01 (5.95e-01, 9.41e-01) | 7.96e-01 (6.04e-01, 8.45e-01) | 7.96e-01 (6.02e-01, 9.22e-01) | 7.96e-01 (5.94e-01, 9.32e-01) | 7.96e-01 (7.96e-01, 2.73e+00) | 7.96e-01 (6.14e-01, 9.37e-01) |
| $\sigma_V$ | 9.05e-01 (6.76e-01, 1.09e+00) | 9.13e-01 (7.06e-01, 1.08e+00) | 9.13e-01 (7.82e-01, 1.00e+00) | 9.14e-01 (6.58e-01, 1.08e+00) | 9.14e-01 (7.16e-01, 1.08e+00) | 9.13e-01 (9.13e-01, 2.31e+00) | 9.14e-01 (6.86e-01, 1.11e+00) |
| SARS-CoV-2 |  |  |  |  |  |  |  |
| $\Delta AIC_c$ | 1.00e+01 | 1.40e+01 | 1.10e+01 | 0.00e+00 | 1.40e+01 | 1.50e+01 | 1.40e+01 |
| $R_0$ | 1.00e+00 (9.93e-01, 2.52e+00) | 1.03e+00 (1.00e+00, 1.89e+01) | 1.01e+00 (9.93e-01, 6.71e+00) | 7.73e+01 (2.20e+00, 1.63e+02) | 1.01e+00 (9.87e-01, 2.70e+00) | 1.02e+00 (5.38e-03, 1.02e+00) | 1.01e+00 (1.01e+00, 2.25e+01) |
| $V_0$ | 3.41e+03 (3.34e+03, 9.73e+05) | 4.43e+04 (5.18e+01, 9.62e+05) | 4.23e+03 (4.23e+03, 9.87e+05) | 1.00e+06 (7.91e+03, 1.00e+06) | 3.54e+03 (8.16e+02, 9.64e+05) | 4.62e+04 (4.62e+04, 6.32e+05) | 3.12e+03 (2.49e+02, 8.62e+05) |
| $\nu_F$ | - | - | - | 1.00e-04 (3.16e-05, 1.00e-04) | 4.60e-06 (8.69e-07, 9.88e-05) | 1.62e-05 (1.62e-05, 6.48e-05) | 1.00e-07 (1.00e-07, 8.68e-03) |
| $\nu_N$ | - | - | - | - | - | 1.69e+04 (1.69e+04, 8.88e+04) | - |
| $\nu_T$ | - | - | 3.31e-08 (3.31e-08, 7.37e+00) | - | - | - | - |
| $\delta_E$ | - | 1.25e+09 (6.84e+08, 9.48e+09) | - | - | - | - | - |
| $\delta_F$ | - | - | - | 3.21e+00 (3.21e+00, 5.57e+03) | 1.92e+04 (4.23e+01, 9.79e+05) | 5.62e+04 (5.62e+04, 7.48e+05) | 1.01e+05 (1.01e+05, 9.56e+05) |
| $\delta_N$ | - | - | - | - | - | 2.40e+03 (2.40e+03, 7.48e+04) | - |
| $\delta_P$ | 1.24e+00 (1.24e+00, 9.64e+05) | 8.94e+01 (8.94e+01, 9.89e+05) | 1.24e+00 (1.24e+00, 9.46e+05) | 2.70e+03 (2.70e+03, 8.29e+05) | 1.00e+00 (1.00e+00, 9.43e+05) | 1.25e+00 (1.25e+00, 3.86e+05) | 1.00e+00 (1.00e+00, 9.52e+05) |
| $\pi$ | - | - | - | - | 9.03e-01 (6.18e-03, 9.87e-01) | - | - |
| $\phi$ | - | - | - | 1.41e+00 (1.41e+00, 5.28e+01) | - | - | - |
| $\psi$ | - | - | - | - | - | 8.81e+04 (8.81e+04, 9.13e+04) | - |
| $\zeta$ | - | - | - | - | - | - | 1.00e+05 (4.47e+01, 1.00e+05) |
| $\rho$ | - | - | - | 1.22e+00 (7.28e-01, 1.38e+00) | - | - | - |
| $\iota_P$ | 1.99e-05 (1.99e-05, 9.25e-01) | 4.82e-04 (4.82e-04, 9.56e-01) | 3.74e-05 (3.74e-05, 9.32e-01) | 1.00e+00 (1.12e-01, 1.00e+00) | 4.38e-05 (4.38e-05, 9.61e-01) | 7.56e-06 (7.56e-06, 7.09e-02) | 4.76e-05 (4.76e-05, 9.72e-01) |
| $\iota_V$ | 9.98e-01 (1.08e-01, 9.98e-01) | 9.96e-01 (1.35e-02, 9.99e-01) | 9.99e-01 (9.65e-03, 9.99e-01) | 1.00e+00 (3.94e-01, 1.00e+00) | 9.91e-01 (2.84e-01, 9.99e-01) | 9.90e-01 (7.85e-01, 9.90e-01) | 9.36e-01 (1.05e-01, 9.96e-01) |
| $\sigma_P$ | 1.63e+00 (1.11e+00, 2.03e+00) | 1.66e+00 (1.25e+00, 2.02e+00) | 1.63e+00 (1.15e+00, 2.01e+00) | 1.98e+00 (1.47e+00, 2.51e+00) | 1.66e+00 (1.12e+00, 2.11e+00) | 1.63e+00 (1.63e+00, 1.69e+00) | 1.66e+00 (1.16e+00, 2.04e+00) |
| $\sigma_V$ | 2.02e+00 (1.29e+00, 2.44e+00) | 2.09e+00 (1.37e+00, 2.56e+00) | 2.02e+00 (1.33e+00, 2.61e+00) | 1.24e+00 (1.06e+00, 1.64e+00) | 2.07e+00 (1.31e+00, 2.43e+00) | 2.02e+00 (2.02e+00, 2.34e+00) | 2.05e+00 (1.41e+00, 2.50e+00) |

## Notes

### Competing Interest Statement

The authors have declared no competing interest.

